# p53 activity links chromosomal instability to pro-migratory extracellular vesicles

**DOI:** 10.64898/2026.10.06.757210

**Authors:** Ruifang Tian, Siqi Zheng, Lin Zhou, Marsudi Siburian, René Wardenaar, H. Rudolf de Boer, Petra L. Bakker, Daniëlle Koot, Xiao Ling, Marcel A.T.M. van Vugt, Floris Foijer

## Abstract

Chromosomal instability (CIN) promotes tumor evolution and metastasis while imposing cellular stress that can limit cell proliferation via activation of p53 and its downstream effector p21. However, how cells recover from this p53-mediated proliferation block while adopting a more malignant fate remains poorly understood. Here, using a p21–RFP reporter in p53-proficient NIH/3T3 cells, we show that acute CIN gives rise to two phenotypically distinct p21 cell populations with divergent recovery trajectories. p21-RFP^LOW^ cells re-enter the cell cycle following a short cell-cycle arrest, while p21-RFP^HIGH^ cells display a more persistent arrest that coincides with a senescence-like state. Longitudinal transcriptomic and extracellular vesicle (EV) proteomic analyses reveal that extracellular-matrix (ECM)- and adhesion-associated programs are initially suppressed following CIN but specifically reactivated in p21-RFP^HIGH^ cells. The content of late-stage p21-RFP^HIGH^-derived EVs is enriched for ECM-, adhesion-, and cytoskeleton-associated proteins compared with that of p21-RFP^LOW^-derived EVs. Consequently, EVs derived from p21-RFP^HIGH^ cells confer increased migration. Together, these findings reveal that CIN can yield distinct p21-expressing cell populations that display different cell-cycle recovery trajectories, EV makeup, and migration-associated paracrine activity.

## Introduction

Chromosomal instability (CIN), defined as an increased rate of alterations in chromosome number or structure, is a hallmark of cancer and correlates with poor prognosis^1–4^. Chromosome mis-segregation can activate p53-dependent surveillance programs, including cell-cycle arrest, senescence, and apoptosis^5–9^. Importantly, these responses are heterogeneous rather than uniform: whereas some cells continue to proliferate after individual chromosome mis-segregation events, others mount a robust p53 response and undergo prolonged arrest, particularly when chromosomal imbalance or associated cellular stresses exceed tolerable levels^5,9,10^. However, how individual p53-proficient cells resolve these different outcomes remains incompletely understood.

The dynamics of the p53–p21 axis are key determinants of cell fate following cellular stress^11,12^. While *TP53* inactivation provides an established route by which cancer cells evade CIN-induced growth suppression^6,8,9^, considerably less is known about how cells that retain an intact p53 response respond to chromosome mis-segregation. In particular, relatively little is known about the role of p21, a downstream effector of p53, in this response. A better understanding of this response is important because cells undergoing prolonged arrest may continue to influence their surroundings despite their limited proliferative activity.

The subset of cells that activates the p53–p21 pathway and undergoes cell-cycle arrest following chromosome mis-segregation may subsequently acquire a senescence-like state and a senescence-associated secretory phenotype (SASP)^5,13^. Senescent and stress-arrested cells can secrete inflammatory cytokines, extracellular-matrix (ECM)-modifying factors, and other bioactive molecules that alter neighboring cells and the surrounding microenvironment^14–19^. CIN itself can similarly trigger context-dependent inflammatory and secretory responses. In some settings, these responses promote immune recognition and clearance of genomically abnormal cells, whereas in others they support tumor-cell survival, invasion, or metastatic behavior^2,4,5,20–22^. CIN-associated inflammatory signaling can also support cancer-cell survival in specific contexts^23^. However, whether distinct p53–p21 response states induced by CIN develop different paracrine programs remains unclear.

Extracellular vesicles (EVs) are important mediators of stress- and senescence-associated paracrine communication^24–26^. EVs can carry bioactive proteins, nucleic acids, and lipids that modulate recipient-cell behavior^25–28^ and may facilitate the removal of intracellular stress-associated material^29^. We have previously shown that p53-deficient cancer cells with induced CIN release EVs that promote migration and invasion of recipient cancer cells^30^.

To better understand a potential contribution of p53 the cellular response to acute CIN, in this study, we devised a NIH/3T3 p21–RFP reporter system driven by the *Cdkn1a* promoter to longitudinally track p53–p21 responses in non-transformed cells. Upon induction of acute CIN through Mps1 inhibition, we identify p21-RFP^HIGH^ and p21-RFP^LOW^ populations with distinct cell-cycle recovery trajectories. Longitudinal transcriptomic and EV proteomic analyses reveal that, at later time points during recovery, p21-RFP^HIGH^ cells maintain a stronger stress response and show a stronger re-emergence of ECM- and adhesion-associated transcriptional programs. Furthermore, EVs derived from p21-RFP^HIGH^ cells are enriched for ECM-, adhesion-, and cytoskeleton-associated proteins. These EVs promote migration of recipient tumor cells more than EVs derived from p21-RFP^LOW^ cells. Together, our findings identify heterogeneous p21 activation as a regulatory feature in the response of cells to CIN and link growth arrest to coordinated transcriptional and extracellular remodeling with distinct paracrine consequences.

## Results

### Chromosomal instability induces heterogeneous p53–p21 activation followed by divergent cell states

To resolve cell-to-cell heterogeneity in the response to acute CIN, we established a p21 reporter system in p53-proficient NIH/3T3 cells, in which TurboRFP expression is driven by the *Cdkn1a* promoter^31^. The reporter enabled the isolation and longitudinal analysis of cells with different p21 response amplitudes (**Fig. 1a**). Acute CIN was induced by transient inhibition of Mps1 with reversine (REV)^5,8,32^. REV treatment shortened mitotic duration and induced frequent chromosome-segregation errors (**Fig. 1b; Supplementary Fig. 1a, b**). Single-cell whole-genome sequencing confirmed widespread and heterogeneous chromosome copy-number alterations in the REV-treated cell populations (**Fig. 1c**). Doxorubicin (DOXO) was included as a DNA-damaging reference condition^33,34^ (**Fig. 1b, c**), and did not result in extensive copy-number alterations. We conclude that REV, but not DOXO, induces extensive karyotypic heterogeneity.

**Figure 1.**
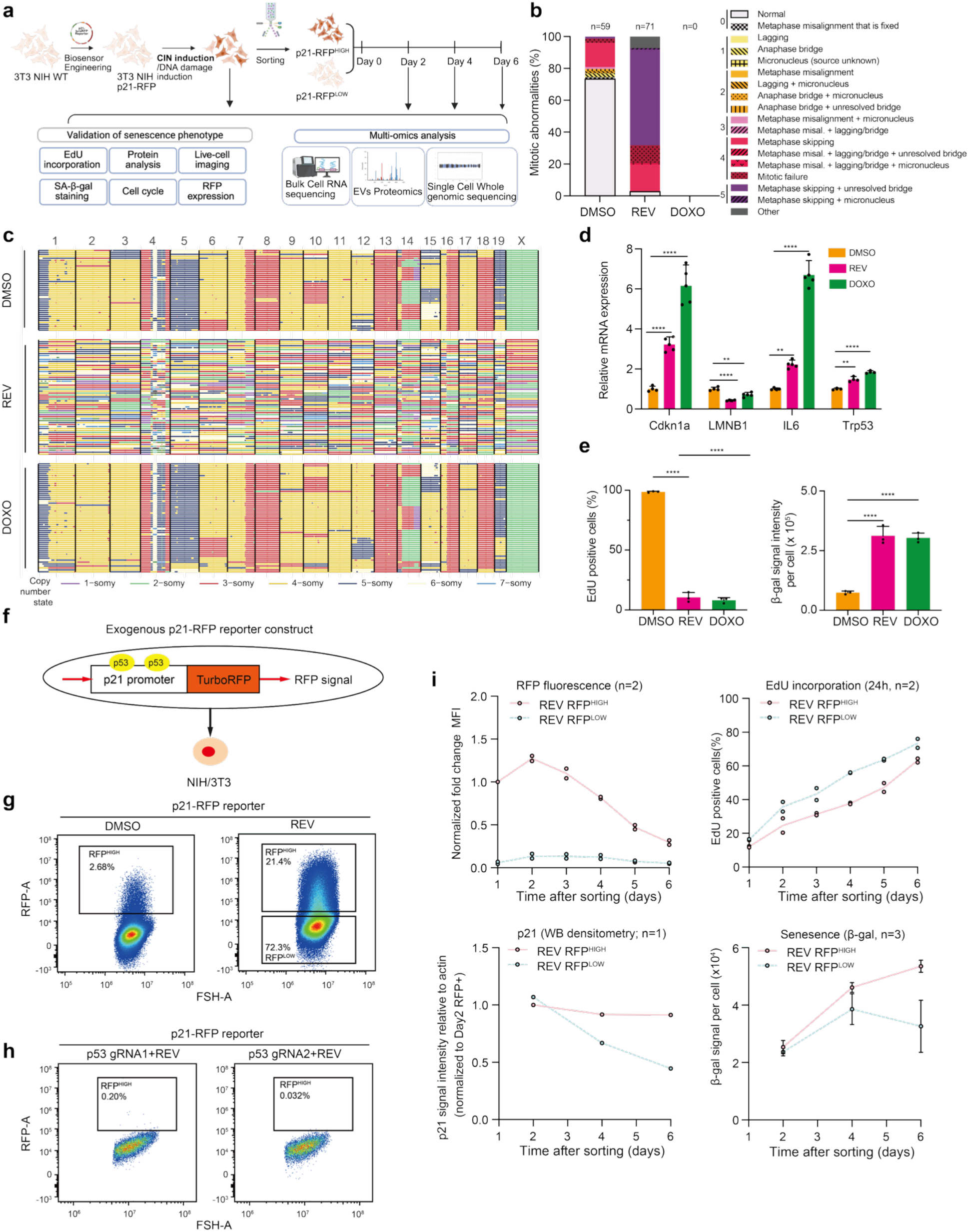
Acute CIN induces heterogeneous p53–p21 responses and distinct but reversible cell-cycle arrest trajectories. **a.** Schematic overview of the experimental design**. b.** Quantification of mitotic errors in NIH/3T3 cells treated with DMSO, reversine (REV; 75 nM), or doxorubicin (DOXO; 250 nM). Mps1 inhibition by REV compromised spindle-assembly-checkpoint control and increased chromosome-segregation errors. No mitotic events were observed in the DOXO-treated group during imaging; therefore, the frequency of mitotic errors could not be calculated for this condition. **c.** Single-cell whole-genome sequencing (scWGS) analysis of NIH/3T3 cells treated with DMSO, REV, or DOXO, showing treatment-associated chromosome copy-number alterations. Each row represents one cell, and colors indicate chromosome copy-number states. **d.** Quantitative PCR analysis of the relative expression of *Trp53*, *Cdkn1a*, Il6, and Lmnb1 in NIH/3T3 cells treated with DMSO, REV, or DOXO. **e.** Quantification of the percentage of EdU-positive cells and SA-β-galactosidase activity following DMSO, REV, or DOXO treatment. SA-β-galactosidase activity is expressed as integrated density per cell. **f.** Schematic of the p21–RFP reporter construct, in which TurboRFP expression is driven by the murine *Cdkn1a* promoter containing p53-binding sites. **g.** Representative flow-cytometry analysis of RFP expression in NIH/3T3 p21–RFP reporter cells treated with DMSO or REV (75 nM, 96 h), showing reporter-defined p21-RFP^HIGH^ and p21-RFP^LOW^ populations. **h.** Validation of reporter dependence on p53 using two independently generated *Trp53*-knockout NIH/3T3 reporter clones analyzed by flow cytometry. **i.** Longitudinal characterization of REV-induced p21-RFP^HIGH^ and p21-RFP^LOW^ populations after sorting. RFP fluorescence and EdU incorporation following a 24h pulse was measured on days 1–6 (n = 2 each). p21 protein abundance was assessed by western blot densitometry (n = 1), and SA-β-galactosidase activity was quantified (n = 3) on days 2, 4, and 6. n denotes independent experiments. Data in panels d and e are presented as mean ± SD. SA-β-galactosidase data in panel i are presented as mean ± SD; RFP and EdU plots show the mean and individual values from two independent experiments. Time-course data are presented descriptively without statistical hypothesis testing. Statistical significance was determined using one-way analysis of variance (ANOVA) followed by Tukey’s multiple comparisons test for panels (**d**) and (**e**) (n = 4 independent experiments for panel **d**; n = 3 independent experiments for panel **e**). Abbreviations: DMSO, vehicle control; REV, reversine; DOXO, doxorubicin. Statistical significance is indicated as follows: *p < 0.05; **p < 0.01; ***p < 0.001; ****p < 0.0001.

At the population level, REV treatment increased p53 and p21 abundance, reduced EdU incorporation, and increased SA-β-galactosidase activity. This was also true for DOXO treatment, which typically involved a larger effect size (**Fig. 1d, e; Supplementary Fig. 1c, d**). To determine whether individual cells activated p21 to different degrees, we next examined p21 reporter activity (**Fig. 1f**). Intriguingly, REV induced heterogeneous p21-RFP fluorescence, allowing separation into p21-RFP^HIGH^ and p21-RFP^LOW^ populations, while the majority of DOXO-treated cells displayed high fluorescence intensity (**Fig. 1g; Supplementary Fig. 1f, g**). Reporter induction was strongly attenuated in two independently generated p53-deficient populations, confirming that p21 expression following CIN is indeed dependent on activation of p53 (**Fig. 1h; Supplementary Fig. 2a, b**). However, loss of p53 did not completely restore proliferation or abolish senescence-associated features (**Supplementary Fig. 2c**). Additional stress-response pathways may therefore contribute to the consequences of acute CIN.

We next asked whether the REV-induced reporter states represented quantitatively different versions of the same response or biologically distinct cell states. Further characterization showed that REV-induced p21-RFP^HIGH^ cells displayed stronger p53–p21 activation and cell-cycle arrest, but also that p21-RFP^LOW^ cells still had a detectable p53–p21 response (**Supplementary Fig. 2d–f**). Within the DOXO-treated population, both populations of cells remained strongly arrested. The DOXO treatment-derived p21-RFP^HIGH^ population was used as a DNA-damage reference in subsequent experiments (**Supplementary Fig. 2g–i**). Thus, reporter heterogeneity following CIN identifies distinct stress-response states rather than simply reflecting stochastic variation in reporter expression.

Longitudinal tracking of the two populations with differential REV-induced p21 levels showed an overall decline in RFP fluorescence and p21 protein levels during recovery. However, p21-RFP^LOW^ cells resumed DNA synthesis substantially earlier than p21-RFP^HIGH^ cells. In line with the reporter state, p21-RFP^HIGH^ cells expressed high levels of p21 for a longer time-period and exhibited delayed recovery of EdU incorporation. The populations also differed in their senescence-associated features. SA-β-galactosidase activity progressively increased in p21-RFP^HIGH^ cells but declined after day 4 in p21-RFP^LOW^ cells during recovery (**Fig. 1i; Supplementary Fig. 3a, b**). Importantly, EdU-positive cells ultimately re-emerged in both populations, indicating that even the sustained p21-RFP^HIGH^ state following acute CIN was not uniformly irreversible. p21-RFP^HIGH^ DOXO treatment-treated cells, on the other hand, maintained high RFP fluorescence and high p21 expression in the absence of DNA synthesis and with increasing SA-β-galactosidase activity at all assessed time points (**Supplementary Fig. 3c–e**).

Together, these findings show that acute CIN induces a heterogeneous p53–p21 response that resolves into distinct, yet partially reversible, cell-state trajectories. Cells mounting a relatively weak p21 response rapidly re-enter the proliferative pool, whereas cells with strong p21 activation remain arrested for longer and progressively acquire senescence-like features. Thus, the magnitude of the early p21 response predicts how individual cells resolve CIN-induced stress.

### Longitudinal transcriptomics reveals distinct cell-cycle recovery and stress-remodeling trajectories following acute CIN

Having established that acute CIN induces p21-RFP^HIGH^ and p21-RFP^LOW^ populations with distinct proliferative recovery kinetics, we next asked whether these phenotypic states were underpinned by divergent molecular programs. To this end, we performed longitudinal RNA sequencing of REV-induced p21-RFP^HIGH^ and p21-RFP^LOW^ cells, together with DOXO-induced p21-RFP^HIGH^ cells on days 2, 4, and 6. Unsorted DMSO-, REV-, and DOXO-treated cells were included as reference populations. Global principal-component analysis (PCA) separated samples according to treatment, reporter state, and time, with DMSO-, REV-, and DOXO-treated populations included as reference conditions (**Fig. 2a**). The experimental groups therefore displayed distinct transcriptional states.

**Figure 2.**
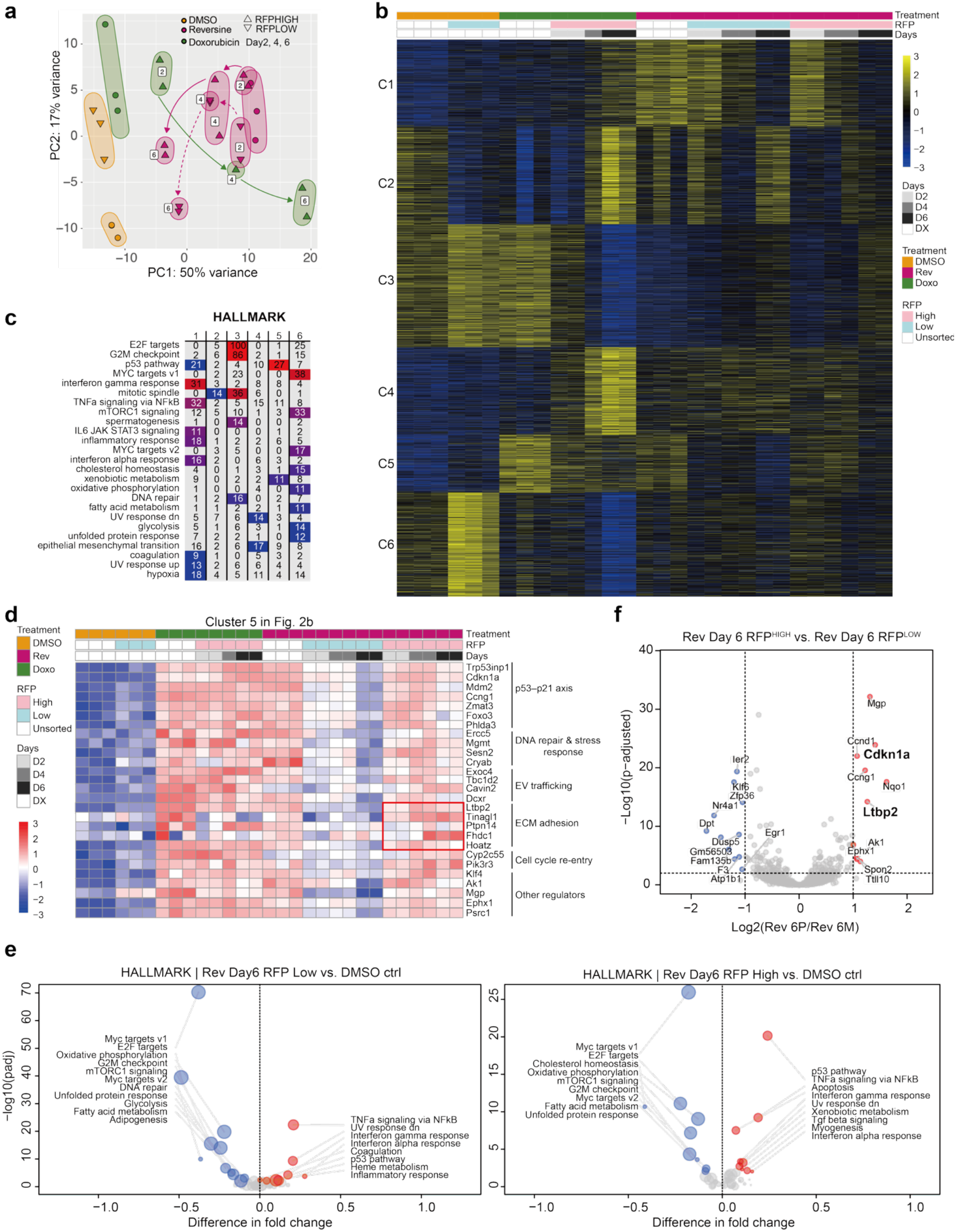
Longitudinal transcriptomics reveals distinct cell-cycle recovery and stress-remodeling trajectories following acute CIN. **a.** Principal component analysis (PCA) of transcriptomes across all 13 experimental groups, including unsorted controls and longitudinally followed reporter-sorted populations. b. K-means clustering analysis of differentially expressed genes (DEGs) across all treatment groups and time points, identifying six major transcriptional clusters. **c.** Hallmark pathway enrichment analysis of the six transcriptional clusters identified in panel b. **d.** Heatmap showing genes upregulated in cluster 5 across treatments and time points, grouped by functional categories. **e.** Hallmark pathway signature analyses comparing REV-induced p21-RFPLOW cells (left) and p21-RFPHIGH cells (right) with DMSO controls at day 6 after sorting. **f.** Volcano plot showing differentially expressed genes between REV-induced p21-RFPHIGH and p21-RFPLOW cells at day 6 after sorting. Positive log₂ fold changes indicate higher expression in p21-RFPHIGH cells, whereas negative values indicate higher expression in p21-RFPLOW cells. Selected genes are labeled.

Analysis of the unsorted populations revealed that REV and DOXO elicited qualitatively different stress responses despite a common p53 activation and cell-cycle suppression. REV treatment resulted in altered inflammatory, cytokine, cytoskeletal, and vesicle-transport programs, whereas DOXO primarily yielded a DNA-damage and DNA repair checkpoint-associated response (**Fig. 2b; Supplementary Fig. 4a–f**). Within the REV-treated population, transcriptomes of p21-RFP^HIGH^ and p21-RFP^LOW^ cells diverged over time, while the transcriptome of DOXO-induced p21-RFP^HIGH^ cells remained transcriptionally distinct from both CIN-induced transcriptional states (**Fig. 2a**). Thus, heterogeneity in the early p21 response to CIN was associated with the emergence of distinct molecular recovery trajectories.

To resolve the temporal programs underlying this divergence, we clustered genes displaying the greatest changes relative to DMSO controls. This identified six transcriptional modules with distinct treatment- and time-dependent dynamics (**Fig. 2b; Supplementary Table 1**). Functional annotation linked these modules to inflammatory and stress signaling (cluster 1), cell-cycle and DNA repair regulation (clusters 2-3), cytoskeletal remodeling (cluster 4), p53-associated arrest (cluster 5), and metabolic and biosynthetic reprogramming (cluster 6) (**Fig. 2c; Supplementary Fig. 5a; Supplementary Table 2**). An inflammatory and stress-response module was rapidly induced after REV treatment but subsequently declined, whereas cell-cycle and DNA repair modules changed dynamically as cells recovered. Consistent with their earlier restoration of DNA synthesis, p21-RFP^LOW^ cells showed stronger recovery of proliferation-associated clusters. By contrast, p21-RFP^HIGH^ cells retained stronger arrest- and stress-associated transcriptional signatures. DOXO-treated cells followed a separate trajectory characterized by sustained DNA-damage signaling (**Fig. 2b, c; Supplementary Fig. 5a**).

The molecular differences between the two CIN-induced populations became particularly clear during the later time points. Genes within cluster 5 were preferentially maintained or reactivated in p21-RFP^HIGH^ cells and encompassed several functionally related programs (**Fig. 2b, d**). These included canonical p53–p21 response genes (*Cdkn1a, Trp53inp1, Mdm2, Ccng1, Zmat3,* and *Phlda3*), DNA-damage and oxidative-stress genes (*Ercc5, Mgmt, Sesn2,* and *Cryab*), and genes implicated in vesicle trafficking (*Exoc4, Tbc1d2, Cavin2,* and *Dcxr*). At later time points during recovery, p21-RFP^HIGH^ cells also showed increased expression of ECM and adhesion-associated genes, including *Ltbp2, Tinagl1, Ptpn14, Fhdc1,* and *Hoatz* (**Fig. 2d**). These observations suggested that persistent p21 activation following CIN was accompanied not only by sustained intracellular stress signaling but also by progressive remodeling of pathways involved in extracellular interactions.

Gene-set enrichment analysis at day 6 further supported the emergence of distinct transcriptional states for p21-RFP^LOW^ and p21-RFP^HIGH^ cells. Although p21-RFP^HIGH^ and p21-RFP^LOW^ populations remained depleted for MYC targets, E2F targets, G2/M checkpoint, DNA repair, and metabolic programs relative to DMSO controls, p21-RFP^HIGH^ cells showed a strong enrichment for p53-, inflammatory- and TGF-β-associated gene sets (**Fig. 2e; Supplementary Fig. 5b**). Direct comparison of the two CIN-induced populations further identified *Cdkn1a, Ccng1, Nqo1, Mgp, Ltbp2, Ak1, Ephx1*, and *Ttll10* among the genes enriched in p21-RFP^HIGH^ cells, whereas *Ier3, Klf9, Nr4a1, Dpt,* and *Dusp5* were more highly expressed in p21-RFP^LOW^ cells (**Fig. 2f**). *Ltbp2*, a fibrillin-associated extracellular matrix gene, was among the most significantly enriched genes in p21-RFP^HIGH^ cells, further supporting the emergence of an ECM-associated state alongside enhanced TGF-β pathway activity during late recovery^35,36^. Its induction, together with enrichment of TGF-β-associated gene sets, suggested extracellular remodeling during prolonged cell-cycle arrest following CIN.

As a complementary approach, we inferred transcription factor activity from our transcriptome data. DOXO-treated cells exhibited increased FOXO, STAT, and SOX9 activity compared to DMSO at all assessed time points, whereas these same programs initially increased and then declined in REV-induced p21-RFP^HIGH^ cells over time, coinciding with partial restoration of E2F- and MYC-associated activity (**Supplementary Fig. 6a, b**). Thus, the p21-RFP^HIGH^ CIN state was functionally distinct from the sustained DNA-damage response and underwent progressive transcriptional remodeling during recovery. Together, these analyses show that heterogeneous p21 responses following acute CIN define molecularly distinct recovery programs. p21-RFP^LOW^ cells rapidly restore their proliferative transcriptional activity, whereas p21-RFP^HIGH^ cells remain in a prolonged stress-associated state that progressively engages ECM, adhesion, and vesicle-trafficking programs. These findings suggest that persistent arrest following CIN is not a transcriptionally static endpoint, but a dynamically remodeling state with the potential to alter extracellular communication.

### Acute CIN increases EV release and remodels the EV-associated proteome

Longitudinal transcriptomics revealed that acute CIN engages vesicle-transport and extracellular-remodeling programs during recovery. We therefore asked whether these intracellular changes were accompanied by an altered extracellular vesicle (EV) output. Building on our previous observation that CIN-exposed breast cancer cells release migration-promoting EVs enriched in EFEMP1^30^, we first examined the population-level EV response to acute CIN in p53-proficient NIH/3T3 p21 reporter cells. EVs were isolated from conditioned media of unsorted DMSO-, REV-, and DOXO-treated cells and analyzed for particle abundance and protein composition, providing a reference for subsequent comparison of p21-reporter populations (**Fig. 3a**). After normalization to viable donor-cell number, treatment with either REV or DOXO significantly increased EV release. Particle-size distributions were similar across groups, with most particles measuring 40–200 nm (**Fig. 3b, c**). Transmission electron microscopy revealed vesicles with the expected morphology (**Supplementary Fig. 7a)**, while enrichment of CD9 and TSG101 and the absence of detectable Calnexin supported EV enrichment (**Supplementary Fig. 7b**). Label-free quantitative proteomics detected more than 800 proteins in EV-enriched isolates, including CD9, CD81, TSG101, PDCD6IP, and SDCBP, whereas Calnexin and GM130 were not detected (**Supplementary Table 3**). Of these proteins, 73 overlapped with the top 100 EV-associated proteins in ExoCarta^37^ (**Supplementary Fig. 7c**). Gene Ontology (GO) Cellular Component analysis further showed enrichment of terms related to vesicle lumen, cell–substrate junctions, and collagen-containing extracellular matrix (**Supplementary Fig. 7d**). Together, these findings supported enrichment of EV populations with small-EV-like characteristics in accordance with current field guidelines^38^.

**Figure 3.**
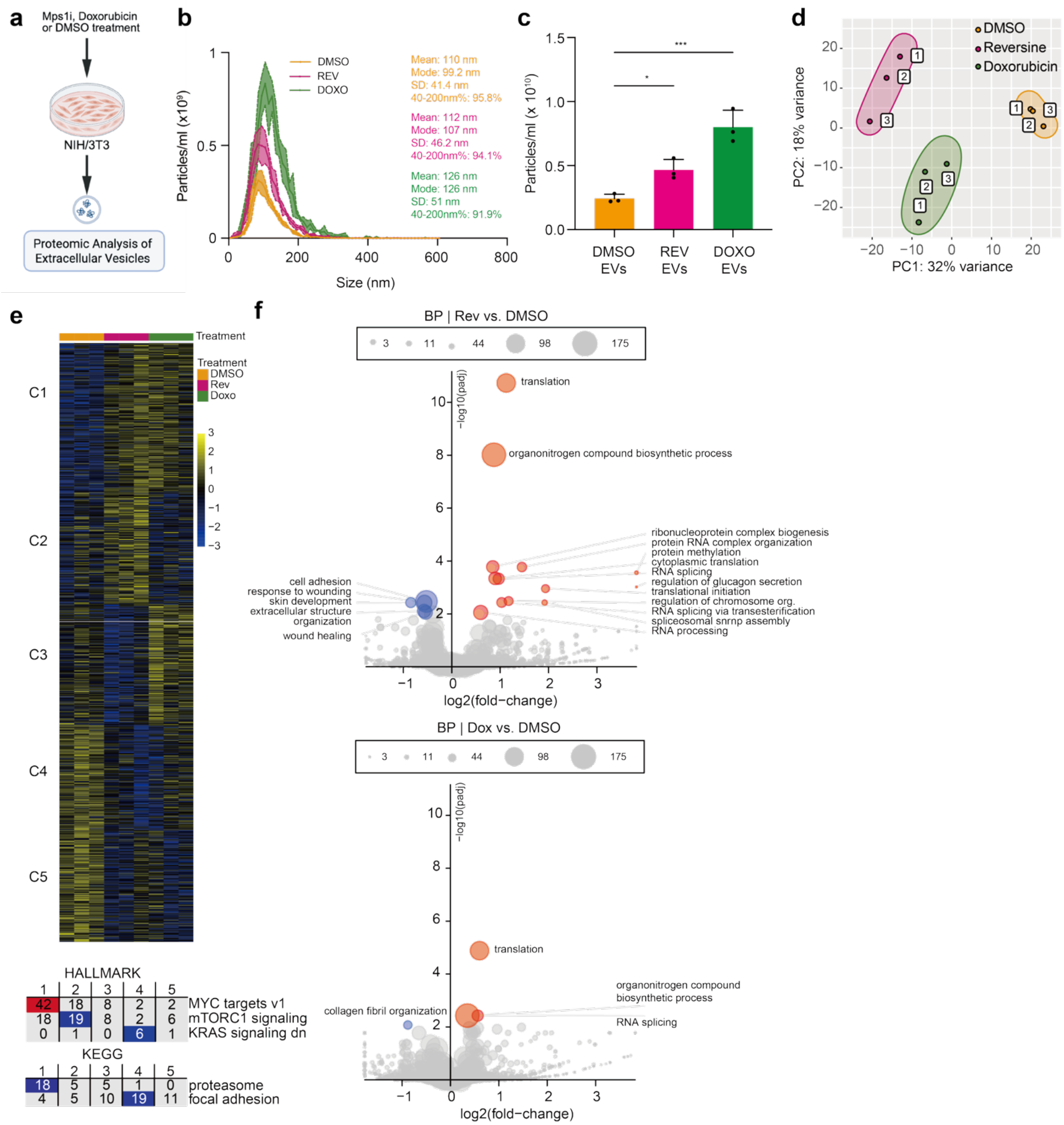
Acute CIN increases EV release and remodels the EV-associated proteome. **a.** Experimental overview of EV isolation and mass spectrometry–based proteomic analysis using unsorted NIH/3T3 p21-RFP reporter cells treated with DMSO, REV, or DOXO. b. Particle-size distributions of EV preparations isolated from conditioned media of DMSO-, REV-, or DOXO-treated cells, measured by nanoparticle tracking analysis (NTA). Lines and shaded areas represent the mean and SD, respectively. c. EV particle numbers measured by NTA and normalized to viable donor-cell number at harvest, expressed per 10⁶ viable donor cells. **d.** Principal component analysis (PCA) of EV proteomes from DMSO-, reversine-, and doxorubicin-treated groups. **e.** K-means clustering of EV-associated proteins across treatment groups, identifying five clusters, with corresponding Hallmark and KEGG pathway enrichment analyses. **f.** GOBP enrichment analysis comparing EVs isolated from reversine- and doxorubicin-treated cells to EVs isolated from DMSO controls. Shown are the mean ± SD values. Statistical significance was determined using one-way analysis of variance (ANOVA) followed by Tukey’s multiple comparisons test for panels (c) (n = 3 independent experiments for panels c). Abbreviations: DMSO, vehicle control; REV, reversine; DOXO, doxorubicin. Statistical significance is indicated as follows: *p < 0.05; **p < 0.01; ***p < 0.001; ****p < 0.0001.

Having established comparable EV preparations across conditions, we next asked whether acute CIN altered cargo composition. PCA clearly separated EV proteomes derived from DMSO-, REV-, and DOXO-treated cells, suggesting treatment-dependent remodeling of the EV content (**Fig. 3d**). Compared to DMSO, REV and DOXO altered 73 and 40 unique proteins, respectively, while 47 proteins were commonly altered under the latter two conditions (**Supplementary Fig. 7e**). Thus, the EV-mediated response is partly overlapping with the DNA-damage response but also has unique features.

To further resolve shared and condition-specific cargo changes, we performed k-means clustering (**Fig. 3e**). The biggest differences were observed within clusters C2 and C3. Proteins in cluster C2 were less abundant in EVs derived from DOXO-treated cells than in EVs isolated from REV-treated cells and included proteins associated with MYC targets and mTORC1-related proteins. In contrast, proteins in cluster C3 were less abundant in EVs isolated from REV-treated cells than in EVs isolated from both DMSO- and DOXO-treated cells and included multiple focal-adhesion-associated proteins (**Fig. 3e**). At the pathway level, compared to EVs isolated from DMSO-treated cells, EVs isolated from REV-treated cells showed an enrichment for proteins involved in translation, RNA processing, ribonucleoprotein-complex organization, and protein transport, accompanied by reduced representation of ECM, cell-adhesion, extracellular-structure, and wound-response proteins. Treatment with DOXO induced a partially overlapping pattern, with increases in RNA processing and translation-associated proteins and decreases in ECM-associated proteins (**Fig. 3f; Supplementary Fig. 8a, b**). Together, these results show that acute CIN increases EV release from non-transformed fibroblasts and leads to an altered protein cargo that is partly shared with EVs secreted by cells with DNA damage, but also partly specific to CIN. Together with the transcriptional changes observed during recovery, these results indicate that acute CIN leads to modulation of intracellular stress programs and extracellular vesicle-mediated communication.

### Longitudinal EV proteomics reveals cell-state-specific cargo remodeling following acute CIN

While our results indicate that acute CIN leads to altered EV release and composition at the population level, whether these changes evolve differently in p21-RFP^LOW^ vs p21-RFP^HIGH^ cells remained unclear. We therefore used the p21 reporter to isolate REV-induced p21-RFP^HIGH^ and p21-RFP^LOW^ cells and collected EVs on days 2, 4, and 6 for quantitative proteomic analysis. EVs from DOXO-induced p21-RFP^HIGH^ cells at matched time points were analyzed in parallel as a DNA damage control. CD9 and TSG101 were detected in the preparations, whereas Calnexin was not detected, supporting EV enrichment (**Supplementary Fig. 9a**).

PCA showed that the EV-associated proteomes derived from the two REV populations displayed distinct temporal patterns that differed from those of the DOXO-derived proteome (**Fig. 4a**), indicating that transcriptional divergence between p21 states was accompanied by time-dependent changes in EV composition. K-means clustering of the complete EV proteomic dataset identified two major clusters (**Fig. 4b**). Cluster 1 was enriched for ECM-, focal-adhesion-, and cell-adhesion-associated proteins. Their abundance initially decreased and subsequently recovered in REV-induced p21-RFP^HIGH^ EVs but declined over time in REV-induced p21-RFP^LOW^ and DOXO-derived EVs when compared to DMSO (**Fig. 4b–d; Supplementary Fig. 9b**). Cluster 2 was enriched for proteins associated with translation, proteostasis, and biosynthesis, including components of ribosomes, proteasomes, and the spliceosome, as well as proteins encoded by MYC- and E2F-target genes. These proteins displayed increased abundance in EVs isolated from both REV-treated p21-RFP^LOW^ and p21-RFP^HIGH^ populations at early time points, while their levels declined at later time points. In EVs isolated from DOXO-treated cells, cluster 2 protein abundance remained relatively stable at all time points (**Fig. 4b–d; Supplementary Fig. 9b**).

**Figure 4.**
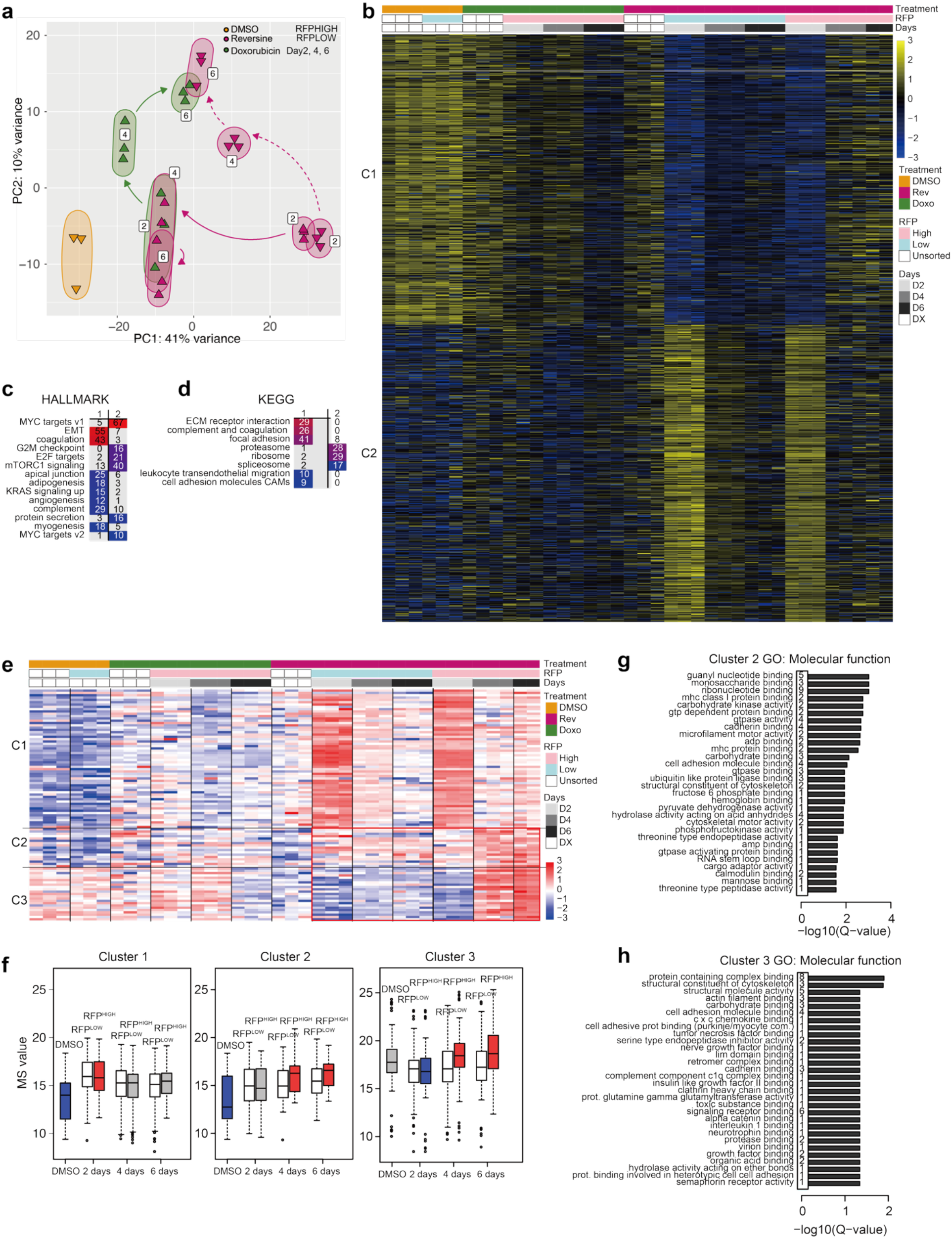
Longitudinal EV proteomics reveals cell-state-specific cargo remodeling following acute CIN. **a.** Principal component analysis (PCA) of EV proteomes derived from sorted cell populations and control. **b.** K-means clustering of EV proteomes from all experimental groups. **c, d**. Hallmark (**c**) and KEGG (**d**) pathway enrichment analyses of the two protein clusters identified in panel **b**. **e.** K-means clustering of proteins significantly increased in day 6 REV-induced p21-RFPHIGH-derived EVs compared with DMSO-derived EVs, identifying three clusters whose abundance patterns are shown across all samples. **f.** Boxplots showing normalized protein intensities (“MS values,” log-transformed and median-normalized) in EVs isolated from reversine-treated p21-RFP^HIGH^ cells over time, compared to proteins in EVs isolated from reversine-treated p21-RFP^LOW^ cells and DMSO control cells. Colors indicate median z-scores corresponding to the heatmap (blue = downregulated, red = upregulated). **g–h**. GO Molecular Function enrichment analyses of proteins in clusters 2 (**g**) and 3 (**h**) identified in panel **e**, using all annotated genes/proteins in the analysis database as the background. Only terms meeting the significance criterion used in the enrichment analysis are shown. Bar lengths indicate −log₁₀(Q-value). Numbers beside the bars indicate the number of proteins from the respective cluster associated with each term.

To characterize the p21-RFP^HIGH^ EV profile at later time points, we selected proteins significantly elevated in EVs isolated from REV-treated p21-RFP^HIGH^ cells at day 6 compared with EVs isolated from DMSO-treated cells and grouped them into three clusters according to their abundance patterns (**Fig. 4e; Supplementary Fig. 9c; Supplementary Table 4**). Proteins in clusters 2 and 3 showed increased abundance at later time points in EVs isolated from p21-RFP^HIGH^ cells compared with EVs isolated from p21-RFP^LOW^ cells and DMSO-treated controls, whereas EV-enriched proteins in cluster 1 showed little difference in abundance between EVs derived from the two REV-induced populations at later time points (**Fig. 4f**). GO Molecular Function analysis linked clusters 2 and 3 to adhesion-, ECM-, and cytoskeleton-associated functions (**Fig. 4g, h; Supplementary Fig. 9d**). Importantly, this phenotype for EVs at later time points mirrored the transcriptional remodeling observed in p21-RFP^HIGH^ cells. Similarly, ECM- and adhesion-associated programs that were initially suppressed following acute CIN were restored during prolonged p21-RFP^HIGH^ recovery at both the RNA and EV-protein levels. In contrast, p21-RFP^LOW^ cells, which resumed proliferation more rapidly, did not show a recovery of these extracellular-remodeling features.

Together, these findings identify a cell-state-specific program that emerges during sustained p21 activation after acute CIN, with p21^-^RFP^HIGH^ cells progressively acquiring an extracellular phenotype characterized by enrichment of ECM-, adhesion-, and cytoskeleton-associated EV cargo.

### p21-RFP^HIGH^-derived EVs are enriched in adhesion-associated proteins and promote scratch-wound closure

To determine whether the altered cargo in p21-RFP^HIGH^ EVs at later time points had functional implications, we first compared the proteomes of EVs derived from p21-RFP^HIGH^ and p21-RFP^LOW^ cells 6 days following sorting (**Fig. 5a**). Proteins enriched in p21-RFP^HIGH^-derived preparations were associated with focal adhesion and extracellular-matrix-related processes (**Fig. 5b**). GO analysis further linked these proteins to cell-adhesion-molecule binding, cytoskeletal interactions and ECM structural function (**Fig. 5c**). Together, these analyses revealed that EVs derived from p21-RFP^HIGH^ cells on day 6 after sorting were enriched for adhesion-, ECM-, and cytoskeleton-associated cargo.

**Figure 5.**
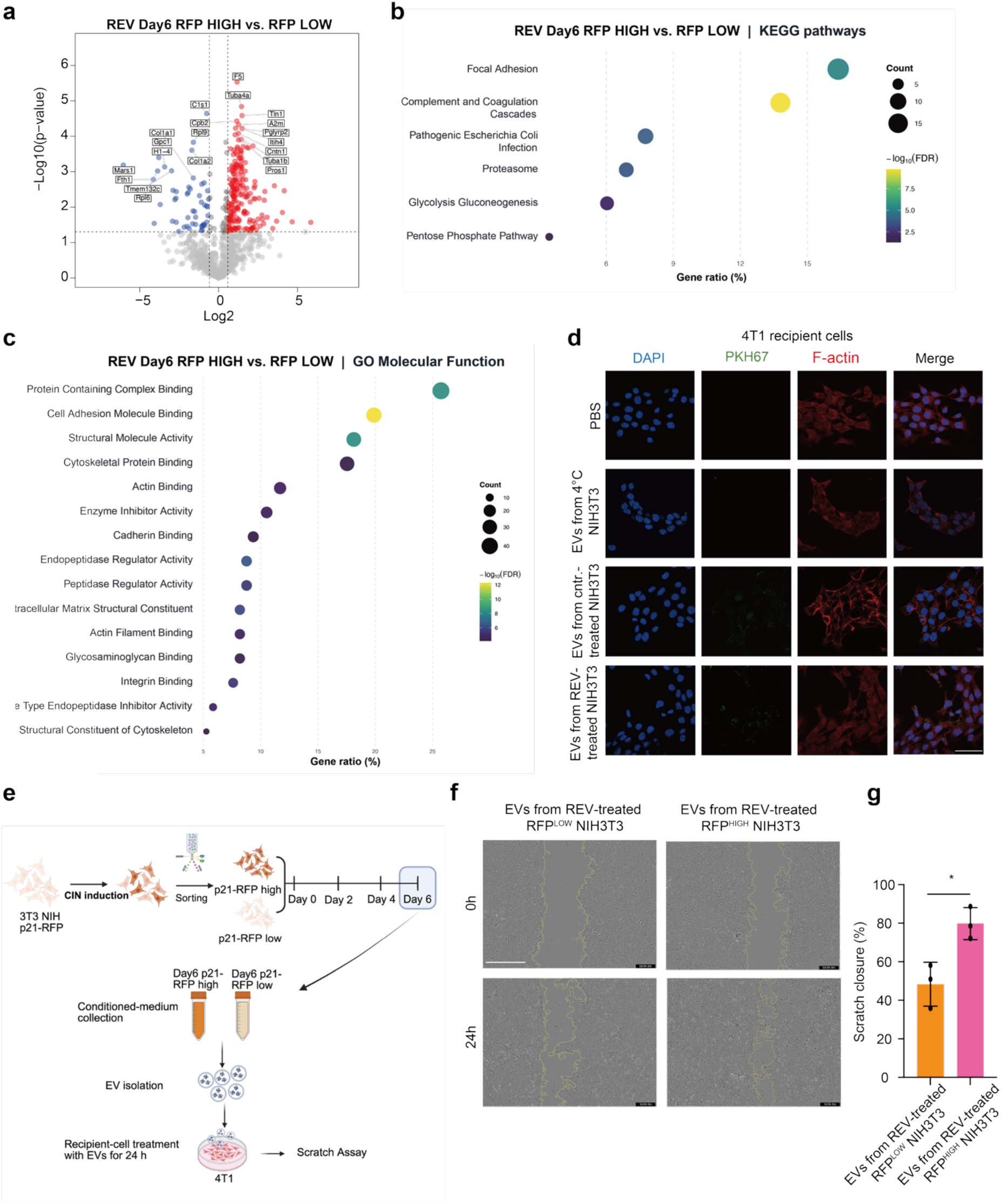
p21-RFP^HIGH^-derived EVs are enriched in adhesion-associated proteins and induce greater scratch-wound closure. **a.** Volcano plot comparing EV protein abundance between day 6 REV-induced p21-RFP^HIGH^ and p21-RFP^LOW^ NIH/3T3 cells. Red and blue indicate significantly increased and decreased proteins, respectively, in p21-RFP^HIGH^ -derived EVs; gray indicates proteins not meeting the significance thresholds. Selected proteins are labeled. **b, c.** KEGG pathway (**b**) and GO Molecular Function (**c**) enrichment analyses of differentially abundant EV proteins identified in the day 6 p21-RFP^HIGH^ and p21-RFP^LOW^ comparison. Bubble size indicates protein count, color represents −log₁₀(FDR), and the x-axis shows the gene ratio (%). **d.** Representative fluorescence images of 4T1 recipient cells incubated for 4 h with PKH67-labeled EV preparations from DMSO- or REV-treated NIH/3T3 cells. PBS and a 4 °C incubation condition were included as controls. PKH67 is shown in green, F-actin in red, and nuclei in blue. Scale bar: 50 μm. **e.** Experimental workflow for EV collection from REV-induced p21-RFP^HIGH^ and p21-RFP^LOW^ NIH/3T3 cells on day 6 after sorting and subsequent treatment of 4T1 recipient cells for scratch-wound assays. **f, g.** Representative images at 0 h (top) and 24 h (bottom) after scratching (**f**) and quantification of scratch-wound closure at 24 h (**g**). Recipient 4T1 cells were treated with EVs derived from REV-induced p21-RFP^LOW^ or p21-RFP^HIGH^ cells. Scale bar: 400 μm. Bars represent mean ± SD, and each point represents one independent experiment (n = 3). Statistical significance was determined using an unpaired two-tailed Student’s t-test for panel **g**. Abbreviations: DMSO, vehicle control; REV, reversine. Statistical significance is indicated as follows: *p < 0.05; **p < 0.01; ***p < 0.001; ****p < 0.0001.

EV-associated ECM and adhesion proteins can influence cell adhesion and directional motility^39^. Recipient 4T1 breast cancer cells were therefore incubated with PKH67-labelled EV preparations from DMSO- or REV-treated NIH/3T3 cells. Cell-associated fluorescence was readily detected at 37°C but was markedly reduced at 4°C, consistent with reduced EV binding to and/or uptake by recipient cells (**Fig. 5d**).

We then compared the functional effects of the EVs isolated from p21-RFP^LOW^ and p21-RFP^HIGH^ cells. As in our proteome measurements, cells were sorted into p21-RFP^HIGH^ and p21-RFP^LOW^ populations after 4 days of REV treatment. The sorted populations were cultured for an additional 6 days, after which EVs were isolated from the corresponding conditioned media. Recipient 4T1 cells were then treated with these EV preparations for 24 h before performing scratch-wound assays (**Fig. 5e**). Assays were performed under serum-free conditions to minimize the contribution of cell proliferation to wound closure. In line with their enrichment for adhesion-, ECM-, and cytoskeleton-associated cargo, EVs derived from p21-RFP^HIGH^ cells induced greater wound closure than EVs derived from p21-RFP^LOW^ cells (**Fig. 5f, g**). Together, these findings demonstrate that p21 expression levels in cells following CIN correlate with distinct EV cargo makeup as well as matched functional consequences, linking sustained cell-cycle arrest as a result of CIN to a pro-migratory paracrine phenotype.

## Discussion

CIN can drive tumor evolution while imposing proliferative and survival stress on individual cells^3,5,7^. The biological consequences of CIN therefore depend not only on whether individual cells continue to divide, but also on how stressed or arrested cells influence their surrounding environment. Increasing evidence is pointing towards an important role for non-cell-autonomous consequences of CIN in cancer cell evolution, including inflammatory signaling and remodeling of the tumor microenvironment^4^. Here, we describe how p21 expression levels following acute CIN correlate with activation of distinct EV programs. Cells with low p21 reporter activity resume DNA synthesis earlier, whereas p21-RFP^HIGH^ cells remain arrested for longer, retain stronger stress-associated signatures, and progressively reactivate ECM- and adhesion-associated transcriptional programs. This is mirrored by changes in the EV proteome and has functional implications, as EVs released by p21-RFP^HIGH^ cells on day 6 after sorting promote greater scratch-wound closure in recipient tumor cells than EVs isolated from p21-RFP^LOW^ cells at the same time point.

Chromosome mis-segregation does not necessarily impose a uniform cell fate. Early studies established p53 as an important barrier to the proliferation of aneuploid cells, whereas later work showed that many cells continue to divide following individual chromosome segregation errors and that p53 activation and subsequent cell-cycle arrest correlate with the severity of the genomic instability ^5,7^. Studies of p53 dynamics and single-cell p21 responses after chemotherapy further demonstrated that response timing and persistence, rather than expression at a single time point, influence recovery and senescence^11,12^. Our findings extend this principle to recovery from acute CIN. Both p21 reporter populations retained detectable p53–p21 activity, yet they differed markedly in the kinetics of DNA-synthesis recovery and acquisition of senescence-associated features. The p21-RFP^HIGH^ and p21-RFP^LOW^ populations should therefore not be interpreted as permanent cell identities, but rather as dynamic states encompassing the magnitude and duration of the response to CIN. However, unlike NIH/3T3 cells arrested by doxorubicin, EdU-positive cells eventually re-emerged in both p21-RFP^LOW^ and p21-RFP^HIGH^ populations, indicating that the arrest induced by CIN in our model system is not uniformly irreversible. This suggests that the CIN-induced p53/p21-mediated cell-cycle arrest is in fact dynamic. In this context, an important next step will be to correlate the severity of genomic damage to p21 reporter activity to determine whether permanent cell-cycle arrest is a possible p53-dependent outcome of CIN.

A central finding of this study is that prolonged arrest following CIN is accompanied by an altered proteomic content of EVs. Indeed, persistent p21 signaling has previously been linked to secretory phenotypes. Sturmlechner et al. showed that p21 can establish an immunosurveillance-associated secretome through Rb-dependent transcriptional mechanisms, whereas persistent DNA damage can induce inflammatory secretion through pathways that are not strictly dependent on p53 or Rb^14,16^. More recently, p21-positive senescent stromal populations have been implicated in immune suppression and tumor progression in vivo^19^. Together, these studies indicate that cell-cycle arrest and secretory activity can be functionally coupled but are not always controlled by the same pathways. Our study builds upon this concept by linking p21 responsiveness following acute CIN to EV composition and paracrine activity. A limitation of our study is that our experiments do not implicate p21 itself as a direct driver of the altered EV composition. Furthermore, our findings suggest that the p53/p21 axis is not solely responsible for the observed phenotypes, as loss of p53 does not fully rescue cell-cycle arrest or the senescence-associated features induced by CIN.

Another important feature of our findings is that the secretory phenotype is subject to temporal changes. This is in line with findings of others that senescence-associated secretory phenotypes are highly dynamic, with their composition varying over time and highly dependent on cellular context ^17,18^. For instance, long-term adaptation to aneuploidy can involve attenuation of CIN-induced stress and inflammatory signaling ^40^. In our model, acute REV treatment initially induced inflammatory and stress-associated transcriptional programs, but these responses declined during recovery. In parallel, p21-RFP^HIGH^ cells selectively restored ECM-, adhesion-, and vesicle-associated transcriptional programs that were suppressed shortly after CIN induction. A similar temporal pattern was observed in the EV-associated proteome. ECM- and adhesion-associated proteins were initially reduced following acute CIN but subsequently recovered or even increased beyond basal levels in EV preparations derived from cells initially sorted as p21-RFP^HIGH^ cells. In contrast, the abundance of these proteins declined over time in EVs isolated from REV-induced p21-RFP^LOW^ cells and DOXO-treated cells. These observations suggest that the extracellular response to CIN differs between individual cells depending on or correlated with p53 activity. These differences are unlikely to reflect cell-cycle arrest alone, as the transcriptional profiles and EV cargo of DOXO-treated cells were clearly distinct from those of CIN-exposed cells.

What can explain these differences? Genome-instability-associated inflammation has been linked to micronuclear activation of cGAS-STING although more recent studies have questioned the role of ruptured micronuclei in the activation of cGAS-STING^41–44^. Conversely, sustained STING signaling as a result of chronic CIN can also contribute to an immunosuppressive tumor microenvironment^4^. These observations emphasize that an inflammatory response can have different biological outcomes depending on the duration and context of genome instability. In our non-transformed cell model, interferon-, NF-κB-, and STAT-associated programs were activated as an early response to CIN but attenuated during recovery when ECM- and adhesion-associated genes were activated. Our data therefore suggest a temporal separation between inflammatory signaling directly following CIN and a later remodeling of the extracellular space via EVs.

This study complements our previous observation that CIN-exposed cancer cells release EVs with migration-promoting activity and altered EFEMP1 content^30^. By segregating donor cells according to their p21 expression and recovery time, our current study reveals that CIN-induced changes to EV cargo are both dynamic and cell-state dependent. REV and DOXO each increased EV release and induced partially overlapping acute changes in EV-associated proteins, indicating that increased EV output may represent a broader stress response. However, the selective re-emergence of ECM-, adhesion-, and cytoskeleton-associated cargo in EVs derived from p21-RFP^HIGH^ cells at later time points during recovery was specific to cells with induced CIN. Particularly, EV composition therefore appears to be highly context-specific.

The altered composition of EVs secreted at later time points by p21-RFP^HIGH^ cells also has functional implications for the microenvironment of the cells, as adhesion-associated EV proteins can influence both interactions with recipient cells and downstream signaling in these cells. For example, EV-associated fibronectin has been implicated in integrin-dependent adhesion and directional motility^27,45^. In our model system, the enrichment of ECM-, integrin-, cadherin-, and cytoskeleton-associated proteins in p21-RFP^HIGH^-derived EV preparations could therefore provide likely candidates for the accelerated wound closure observed in recipient cells. However, the exact mechanism driving increased migration and invasion remains to be resolved. Furthermore, our model system does not take into account interactions with immune cells, which should be a topic of follow-up studies.

Our study primarily uses pharmacologically induced CIN in NIH/3T3 fibroblasts, and the generality of our findings in the context of CIN will require further validation using alternative CIN-induction strategies, additional primary cell models, and models for chronic CIN. Furthermore, the exact mechanisms driving the observed increased cell migration still need to be resolved, for instance by modulating expression levels of candidate proteins whose presence is altered in EVs secreted by p21-RFP^HIGH^ cells at later time points during recovery. Finally, to better understand how these p21-RFP^LOW^ and p21-RFP^HIGH^ populations following CIN will affect innate immunity and potentially metastasis, *in vivo* modelling will be required, which will be key to determining whether this mechanism also provides a therapeutic vulnerability to modulate immune recognition or prevent distant seeding of cancer cells.

Collectively, our findings identify the p53 response through p21 expression following CIN as a feature that correlates with cellular recovery and altered extracellular communication. Cells with lower p21 expression re-enter the proliferative pool, whereas cells with higher p21 expression remain arrested for longer, which coincides with a specific transcriptional program and EV cargo. Cells that maintain high p21 expression following CIN are characterized by restoration of ECM-, adhesion-, and cytoskeleton-associated extracellular programs and by enhanced migration-associated activity toward neighboring tumor cells via EVs (**Fig. 6**). Our findings support a model in which CIN does not render stressed non-transformed cells biologically inert; instead, a subset of persistently arrested cells adopts a distinct extracellular signaling state that may influence neighboring cells during recovery from genome instability.

**Figure 6.**
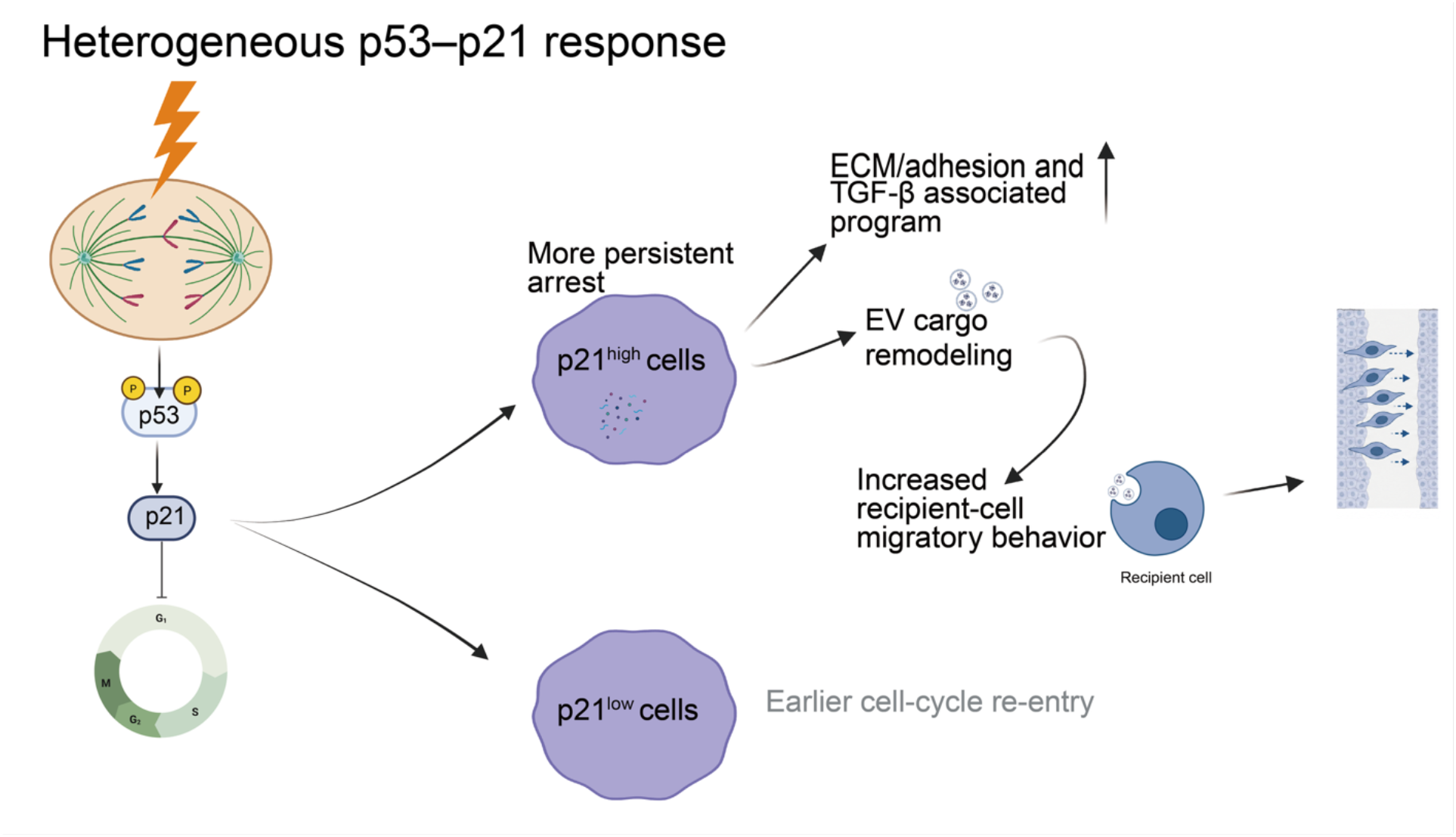
Working model linking heterogeneous p53–p21 responses to cell-cycle recovery, EV remodeling, and paracrine activity following acute CIN. Acute CIN induces heterogeneous p53–p21 responses, generating p21^HIGH^ and p21^LOW^ cell states with distinct recovery trajectories. p21^LOW^ cells re-enter the cell cycle earlier, whereas p21^HIGH^ cells maintain a more persistent arrest and progressively develop ECM-, adhesion-, and TGF-β-associated transcriptional programs. This state is accompanied by remodeling of EV protein cargo and the release of EVs enriched in ECM-, adhesion-, and cytoskeleton-associated proteins. Compared with EVs from p21-RFP^LOW^ cells, EVs released by p21^HIGH^ cells enhance scratch-wound closure in recipient tumor cells, linking persistent p21 activation to a distinct, migration-associated paracrine program.

## Materials and Methods

### Cell culture

NIH/3T3 fibroblast cells and 293FT cells were cultured in Dulbecco’s Modified Eagle Medium (DMEM) (Gibco, 31966-021) with 10% fetal bovine serum (FBS) (Thermo Fisher Scientific, 11573397) and 1% penicillin/streptomycin (P/S) (Gibco, 15140-122). 4T1 cell lines were cultured in RPMI 1640 medium with GlutaMAX supplement (Gibco, 72400-054), 10% FBS, and 1% P/S. All cell lines were obtained from the American Type Culture Collection (ATCC). All cell lines were routinely tested for Mycoplasma contamination and were confirmed to be negative. Cell lines were grown at 37°C and 5% CO2.

### Drug Treatments for Inducing CIN and Senescence

To induce CIN, cells were treated with 75 nM reversine (Sigma, R3904) for 96 hours^20–22^. The reversine dose and treatment duration were selected based on preliminary optimization experiments to achieve cell-cycle arrest and senescence induction while maintaining sufficient cell viability for subsequent sorting and downstream analyses. To generate DNA-damage–induced senescence, cells were exposed to 250 nM doxorubicin (MedChemExpress, HY-15142) for 24 hours, followed by 48 hours in drug-free medium ^23,24^. The dose was based on previous studies, and the recovery period was chosen to assess early DNA-damage–induced cell-cycle arrest prior to sorting and downstream analyses. DMSO (0.1%) (ITW Reagents, A3672,0250) was used as a vehicle control in all experiments.

### Lentiviral Transduction

Lentiviral particles encoding H2B–mCherry or other constructs were produced by transfecting 293FT cells with the corresponding expression plasmid together with the packaging plasmids psPAX2 and pMD2.G (Addgene #12260 and #12259). The H2B–mCherry expression vector was generated by subcloning the H2B–mCherry fragment from an existing construct using the restriction sites NotI and BsrGI (Bioke). At 48–72 hours after transfection, viral supernatants were harvested, cleared through a 0.45-µm filter, and applied to target cells in the presence of polybrene (8 μg/ml) to enhance transduction efficiency. Transduced cells were expanded and used for subsequent experiments.

### Generation of stable p21-RFP reporter cells

NIH/3T3 cells were transfected with the p21 promoter–driven TurboRFP reporter plasmid (pTurboRFP-PRL, Addgene #52432)^31^ using Lipofectamine 3000 (Thermo Fisher). Forty-eight hours after transfection, cells were subjected to G418 selection (400 μg/ml) (Bio-connect, SC-29065A) for two weeks. Surviving colonies were isolated as single clones and expanded. Clones exhibiting robust and reversible RFP induction upon stress treatment were selected for subsequent experiments.

### Genetic Modification Using CRISPR/Cas9

To generate p53 knockout (*Trp53*-KO) NIH/3T3 cells, we employed the CRISPR/Cas9 system using the plasmid pSpCas9(BB)-2A-Puro V2.0 (PX459, Addgene #62988). Two guide RNAs (sgRNAs) targeting exon 5 (5′-TGAGGGCTTACCATCACCAT-3′) and exon 6 (5′-GGTACCTTATGAGCCACCCG-3′) of the *Trp53* gene were designed and cloned into PX459 via BbsI restriction sites. NIH/3T3 cells were transfected separately with 2 µg of either sgRNA plasmid using FuGene® (Promega, E2311) following the manufacturer’s protocol. 48 hours post-transfection, cells were selected with puromycin (2 µg/ml) for 48 hours to enrich for transfected populations. Surviving cells were plated for single-cell cloning, and individual colonies were expanded to establish stable monoclonal lines. Knockout efficiency was validated by Western blotting. To confirm functional loss of p53 activity, selected clones were treated with Nutlin-3, a p53 stabilizer.

### Real-time quantitative PCR

Total RNA was extracted and purified from cultured cells using the RNeasy Mini Kit (Qiagen, 74136). For reverse transcription, 1.5 μg of total RNA was converted into cDNA in a 20 μL reaction mixture containing LunaScript RT SuperMix Kit (NEB, M3010X). Quantitative PCR was performed using iTaq Universal SYBR Green Supermix (Bio-rad, 1725124) on a LightCycler® 480 system (Roche). Gene expression levels were normalized to actin, which served as the internal reference. Primer sequences are listed in **Supplementary Table 5 (Supplementary information)**.

### Time-lapse Imaging

Following previously established procedures^46^, time-lapse imaging was performed on a DeltaVision Elite microscope (Applied Precision Ltd./GE Healthcare) equipped with a CoolSNAP HQ2 camera and 20× 0.75 NA or 40× 0.6 NA immersion objectives. To enable visualization of chromatin dynamics, we introduced the H2B–mCherry construct into the cell lines using lentiviral transduction. A total of 80,000–90,000 H2B–mCherry–expressing cells were seeded in 4-well imaging chambers (LabTech) and cultured overnight. Cells were treated with 75 nM reversine 1 h before imaging, or with 250 nM doxorubicin for 24 h followed by drug washout prior to imaging. Cells were imaged for at least 24 h, and z-stacks of 40 images at 0.5 μm intervals were acquired every 6 min. Image analysis was performed using ICY software (Institut Pasteur), and only cells that entered mitosis and remained in frame throughout the imaging session were included in the analysis.

### EdU incorporation assay

Cell proliferation was evaluated by 5-ethynyl-2′-deoxyuridine (EdU; Sigma-Aldrich, 900584) incorporation using both fluorescence microscopy and flow cytometry. For microscopy, 20,000 NIH/3T3 cells were seeded on glass coverslips in 12-well plates and cultured overnight. Cells were then pulsed with 10 μM EdU for 2 or 24 hours, fixed with 4% formaldehyde solution (diluted from 37–41% formaldehyde stock; Thermo Fisher Scientific, F/1501/PB15) in PBS (Thermo Fisher Scientific, 12037539), and stained using a Click-iT reaction containing 2 mM CuSO₄, 4 μM sulfo-Cy3-azide (Lumiprobe, cat. no. D1330), and 20 mg/ml sodium ascorbate. Fluorescence images were captured on an Olympus microscope, and EdU-positive cells were quantified using ImageJ.

For flow cytometry, cells were labeled with 10 μM EdU for 2 h, fixed, and permeabilized with 0.1% Triton X-100 (Sigma-Aldrich, 9036-19-5) in PBS. The Click-iT reaction was carried out using sulfo-Cy5-azide (Lumiprobe, A3330) to detect EdU incorporation, followed by DAPI (Sigma, D9542) staining for DNA content. Samples were analyzed on a Quanteon flow cytometer (NovoCyte, Agilent), and data were processed using FlowJo software (version 10.8.1; FlowJo LLC).

### SA-β-gal staining

For senescence-associated β-galactosidase (SA-β-gal) staining, cells were seeded in 12-well plates and fixed with 2% formaldehyde (Thermo Fisher Scientific, 28908) and 0.2% glutaraldehyde (Thermo Fisher Scientific, 8.20603.1000) for 10 min at room temperature. Cells were then incubated with freshly prepared staining buffer containing 1 mg/ml X-gal (Cayman Chemical, 7240-90-6) dissolved in dimethylformamide (Merck Millipore, 1.10983.1000), 40 mM citric acid (Sigma-Aldrich, C1909) /sodium phosphate (Thermo Fisher Scientific, 424380010) buffer (pH 6.0), 5 mM potassium ferrocyanide (Thermo Fisher Scientific, 10216460), 5 mM potassium ferricyanide (Thermo Fisher Scientific, 10414583), 150 mM NaCl (Merck Millipore, 1.06404.1000), and 2 mM MgCl₂ (Fisher Chemicals, cat. no. M/0550/53). Staining was performed at 37 °C (without CO₂) for 72 hours. Images were captured at 20× magnification using an Olympus microscope, and the intensity of β-gal-positive cells was quantified using Fiji/ImageJ2^47^ (version 2.16.0; ImageJ version 1.54p; NIH, USA).

### Western blot analysis

Cells or EVs were lysed in RIPA buffer (150 mM NaCl, 50 mM Tris-HCl, pH 8.0, 1% NP-40, 0.5% sodium deoxycholate, and 0.1% SDS) supplemented with a protease inhibitor cocktail (Roche) for 30 min on ice. Protein concentrations were determined using the BCA Protein Assay Kit (Pierce™, Thermo Scientific). Cell lysates were mixed with 5× SDS loading buffer, whereas EV lysates were combined with 4× LDS buffer. Samples were heated at 95 °C for 5 min. Equal amounts of protein (20–50 µg) were resolved on 7.5–10% SDS–PAGE gels and transferred onto PVDF membranes. Membranes were blocked with Odyssey Blocking Buffer (LI-COR Biosciences) or 5% non-fat milk in TBS–0.1% Tween-20 (TBST) and incubated overnight at 4 °C with primary antibodies: Vinculin (mouse, Sigma-Aldrich V9131, 1:20,000), p21 (rabbit, Abcam ab188224, 1:6,000), p53 (rabbit, CST 32532S, 1:1,000), CD9 (rabbit, Invitrogen SA35-08, 1:1,000), TSG101 (mouse, GeneTex GTX70255, 1:1,000), β-Actin (mouse, CST 3700S, 1:1,000), and Calnexin (rabbit, Bio-Techne NB100-1965, 1:1,000). After washing, membranes were incubated with IRDye 800CW Goat anti-Rabbit IgG (H+L) and IRDye 680CW Goat anti-Mouse IgG (H+L) (both 1:15,000; LI-COR) for 1 h at room temperature. Fluorescent signals were detected using the Odyssey Imaging System (LI-COR Biosciences) and quantified with Fiji/ImageJ2 (version 2.16.0; ImageJ version 1.54p; NIH, USA).

### Flow cytometry and cell sorting

Flow cytometry and cell sorting were performed to analyze and isolate p21-RFP^HIGH^ and p21-RFP^LOW^ populations. Cells were harvested using 0.05% trypsin-EDTA, washed twice with cold PBS containing 2% FBS, and filtered through a 40-µm cell strainer to obtain a single-cell suspension. For flow cytometric analysis, samples were analyzed on a Quanteon flow cytometer (NovoCyte, Agilent), and data were processed using FlowJo software (version 10.8.1; FlowJo LLC). For fluorescence-activated cell sorting (FACS), cells were sorted based on RFP fluorescence intensity using a MoFlo Astrios cell sorter (Beckman Coulter). Dead cells and debris were excluded by forward/side scatter and DAPI staining where indicated. The sorted p21-RFP^HIGH^ and p21-RFP^LOW^ populations were collected in complete medium and used for downstream assays, including RNA extraction, immunoblotting, and EV isolation.

### Extracellular Vesicle Isolation

EVs were isolated from the conditioned media of NIH/3T3 cells as previously described^38,48^. To minimize serum-derived contamination, cells were cultured in medium supplemented with 5% EV-depleted FBS (prepared by ultracentrifugation at 100,000 × g for 16 h). When cultures reached 50–60% confluence, the medium was replaced with fresh EV-depleted medium, and cells were incubated for an additional 48–72 h prior to EV collection. For each condition (DMSO, reversine, and doxorubicin treatments), EVs were isolated using a sequential centrifugation and ultrafiltration protocol. Briefly, conditioned media were subjected to stepwise centrifugation at 300 × g for 10 min, 2,000 × g for 20 min, and 10,000 × g for 30 min to remove cells, debris, and large vesicles. The clarified supernatant was subsequently concentrated and purified using Vivaspin® 20 centrifugal filter units (100 kDa MWCO Polyethersulfone, Cytiva, 28-9323-63), followed by a final centrifugation step at 100,000 × g for 70 min to pellet small EVs. Pellets were washed once with sterile PBS, centrifuged again at 100,000 × g for 70 min, and finally resuspended in PBS for downstream analyses.

For sorted RFP^HIGH^ (p21-RFP^HIGH^) and RFP^LOW^ (p21-RFP^LOW^) populations, EVs were collected sequentially over a 6-day period after sorting to capture dynamic changes in vesicle release. Specifically, fresh EV-depleted medium was added immediately after sorting, and conditioned media were harvested every two days (Day 2, Day 4, and Day 6). Each batch was independently processed as described above, and the three time-point EV fractions were analyzed separately by mass spectrometry and functional assays to reflect temporal secretion dynamics.

### Transmission Electron Microscopy (TEM)

For ultrastructural visualization, EVs were fixed in 2% paraformaldehyde prepared in 0.1 M sodium cacodylate buffer (pH 7.4), as described previously^38,48,49^. Carbon-coated Formvar grids were used to provide enhanced film stability. Before EV loading, grids were treated with 1% Alcian blue in 1% acetic acid and rinsed thoroughly with double-distilled water to increase hydrophilicity. A 10 µL aliquot of the EV suspension was placed on the grid and incubated for 30 min to allow vesicle adhesion. Grids were subsequently washed and negatively stained with either 0.5% uranyl acetate or 1% phosphotungstic acid (neutral pH) for 20 min at room temperature. Samples were examined using a Talos 200 (Thermo Fisher Scientific) transmission electron microscope.

### Nanoparticle Tracking Analysis (NTA)

EV size distribution and particle concentration were measured on a ZetaView PMX-130 MONO Laser Nanoparticle Tracking Analyzer (Particle Metrix, Ammersee, Germany). Calibration was performed using 110 nm polystyrene reference beads, and all measurements were conducted at 25 °C. EV samples were diluted in sterile 1× PBS to achieve optimal particle concentration. Each sample was recorded at 11 positions and analyzed using ZetaView software v8.04.02. Every measurement was performed in triplicate for reproducibility. Donor-cell viability was assessed by trypan blue exclusion at harvest. Total recovered particle numbers were calculated from the measured particle concentration, dilution factor, and final EV suspension volume, and normalized to viable donor-cell number.

### Scratch Assays

For scratch assays, cells were detached, counted, and seeded in 6 cm dishes to reach full confluence. Cells were then reseeded into 12-well plates and allowed to attach overnight. Recipient 4T1 cells were incubated with EV preparations for 24 h before scratching (EVs, 10–20 µg/ml total protein). A straight wound was created across the monolayer using a 1 mm sterile pipette tip. After scratching, the wells were gently washed twice with PBS to remove detached cells and debris. Fresh serum-free medium was added immediately after scratching. Scratch closure was monitored using an IncuCyte live-cell imaging system at 10× magnification, capturing images every 2 hours for 24–48 hours. Image analysis and quantification of wound closure were performed using Fiji (version 2.16.0; ImageJ version 1.54p; NIH, USA) software.

### EV Uptake Assay

EVs were labeled with PKH67 (Sigma-Aldrich, MINI67-1KT) following a dye-quench–wash protocol to remove free dye. Briefly, EVs (10–30 µg total protein by BCA) were diluted in 250 µL Diluent C (kit supplied). Separately, 2 µM PKH67 was prepared in 250 µL Diluent C and mixed with the EV suspension for 5 min at room temperature with gentle tapping. Labeling was quenched by adding an equal volume of 1% BSA in PBS for 1–2 min. Labeled EVs were washed twice by ultracentrifugation (100,000 × g, 70 min, 4 °C), resuspended in PBS, and quantified (BCA and/or NTA). A dye-only control (no EVs, processed identically) was included to exclude dye micelles. For uptake, recipient 4T1 cells grown on glass coverslips (24-well plates) were incubated with PKH67-EVs (10–20 µg/ml, in serum-free medium) for 4 h at 37 °C. A 4 °C condition was run in parallel as an energy-dependent uptake negative control^50^. After incubation, cells were washed 3× with PBS, fixed with 4% formaldehyde (15 min), permeabilized with 0.1% Triton X-100 (5–10 min), and blocked with 3% FBS (30 min). F-actin was stained with Phalloidin-iFluor 555 (Abcam, ab176756), nuclei with DAPI (Sigma, D9542). Images were acquired on a Leica SP8 confocal microscope using identical settings across conditions.

### Single-Cell Whole-Genome Sequencing (scWGS)

Cells were harvested for nuclei isolation and stained with Hoechst in nuclear isolation buffer. Single nuclei were sorted into 96-well plates using a FACSJazz cell sorter (BD Biosciences). Single-cell DNA libraries were then prepared and sequenced on an Illumina platform as previously described^51^. Karyotype profiles were analyzed using AneuFinder68 to detect copy-number variations and assess chromosomal instability.

### RNA Sequencing Sample and Library Preparation

RNA quality control (QC) was performed by assessing RNA concentration using a Nanodrop spectrophotometer, followed by evaluation of RNA integrity and RIN values using the High Sensitivity RNA ScreenTape or RNA ScreenTape assay on the Agilent 4200 TapeStation system. Library preparation was carried out using the Smart-3seq protocol on an Agilent Bravo Automated Liquid Handling Platform, with 100 ng of total RNA as input per sample. For low-concentration samples, a pre-bead cleanup step (poly(A) RNA capture) using SPRI beads was included. Libraries were indexed with NEBNext Multiplex Oligos for Illumina (E6440L) and amplified by 12 PCR cycles. Bead cleanup was performed twice at a 0.8× ratio. Post-library QC involved concentration measurement using the Qubit 1X dsDNA HS Assay Kit and BioTek Synergy H1 Multimode Reader, and molarity determination using the High Sensitivity D5000 or D5000 ScreenTape assay on the Agilent 4200 TapeStation.Libraries were diluted to 4 nM, pooled, and verified for concentration and molarity using the Qubit 1X dsDNA HS Assay Kit, Qubit 4 Fluorometer, and HSD5000 ScreenTape assay. Sequencing was performed on the Illumina NextSeq 2000 platform using a P2 100-cycle kit. Run parameters included read 1: 113 bp, read 2: 7 bp, index 1: 9 bp, index 2: 9 bp, with a loaded molarity of 750 pM and 5% PhiX spike-in.

### Proteomics sample preparation

Fifteen micrograms of total protein from extracellular vesicle samples were mixed with LDS loading buffer (Thermo Scientific) in a total volume of 20 μL, incubated at 70°C for 10 minutes, and loaded onto precast 4–12% Bis-Tris gels (Thermo Scientific). Samples were run briefly into the gel (approximately 5 min at 100 V). The gel was stained with rapid Coomassie blue protein stain (Westburg), and after destaining with MilliQ H₂O, the band containing all proteins was excised.

The gel band was sliced into small pieces and washed sequentially with 30% and 50% (v/v) acetonitrile in 100 mM ammonium bicarbonate (each for 30 min at RT with mixing at 500 rpm), followed by 100% acetonitrile for 5 min, before drying at 37°C. Proteins were reduced with 30 μL of 10 mM dithiothreitol (in 100 mM ammonium bicarbonate, 30 min, 55°C) and alkylated with 30 μL of 55 mM iodoacetamide (in 100 mM ammonium bicarbonate, 30 min, RT, in the dark). Gel pieces were then washed with 100% acetonitrile for 30 min (500 rpm) and dried at 37°C, followed by overnight digestion with 30 μL of 10 ng/μL trypsin (sequencing-grade modified trypsin V5111, Promega) at 37°C. The following day, peptides were eluted with 50 μL of 75% (v/v) acetonitrile containing 5% (v/v) formic acid (20 min, RT, 500 rpm). Eluted fractions were dried under vacuum and resuspended in 300 μL of 0.1% (v/v) formic acid.

## Statistical Analysis

Statistical analyses were performed only for experiments with at least three independent biological replicates (n ≥ 3). Data from these experiments are presented as mean ± standard deviation (SD). For comparisons between two groups, statistical significance was assessed using an unpaired two-tailed Student’s t-test. For comparisons involving three or more groups, one-way analysis of variance (ANOVA) followed by Tukey’s multiple comparisons test was applied. Experiments with fewer than three biological replicates (n = 1–2), including time-course measurements and Western blot densitometry, were analyzed descriptively and were not subjected to statistical hypothesis testing, consistent with recommended practices. Time-course plots represent mean values (where applicable) and are intended to illustrate temporal trends rather than statistical comparisons. All statistical analyses were performed using GraphPad Prism 9.5.1. A p value < 0.05 was considered statistically significant (*p < 0.05; **p < 0.01; ***p < 0.001; ****p < 0.0001). No formal method was used to predetermine sample size, and data distribution was assumed to be normal but not formally tested.

## Data availability

All RNA-sequencing, scWGS, and proteomics data generated in this study have been deposited in public repositories. RNA-sequencing data have been deposited in ArrayExpress under accession number E-MTAB-16872. Single-cell whole-genome sequencing (scWGS) data have been deposited in the European Nucleotide Archive (ENA) under accession number PRJEB111512. Proteomics data have been deposited to the ProteomeXchange Consortium via the PRIDE partner repository under accession number PXD076387. All other data supporting this study are provided within the article and its Supplementary Information.

## Acknowledgements

We thank the members of the Foijer lab, the van Vugt lab, and other research groups at ERIBA for valuable discussions and support throughout this work. We are grateful to Johan Teunis and Theo Bijma from the Flow Cytometry Facility for their assistance with cell sorting, and to Nancy Halsema and Laura Kempe at iPsomics for their help with RNA sequencing. We also thank Ben Giepmans, Kim Kats, and Anouk Wolters for their support with TEM imaging. Mass spectrometry analyses were performed at the Interfaculty Mass Spectrometry Center, with expert assistance from Marcel P. de Vries, Hjalmar Permentier, and Dr. Karin Wolters. We further acknowledge Joop de Vries, Willem Woudstra, and Hélder A. Santos for providing access to nanoparticle quantification equipment. Part of this work was carried out at the UMCG Imaging and Microscopy Center (UMIC), supported by the Netherlands Electron Microscopy Infrastructure (NEMI; NWO 184.034.014). Schematic overviews were created using BioRender.com.

Siqi Zheng and Ruifang Tian were supported by personal fellowships from the China Scholarship Council (CSC). Furthermore, this work was supported by an NWO Vici grant (09150182210049) FF. Additional support for Ruifang Tian was provided by the De Cock-Hadders Foundation.

## Author Contributions

RT and SZ designed the study. RT conducted most experiments, data analysis, and figure preparation. SZ performed extracellular vesicle isolation and related experiments. LZ performed and assisted with key molecular and cell-based experiments. RW analyzed the multi-omics datasets and processed proteomics data with support from HRdB. DK performed immunofluorescence experiments, and XL assisted with molecular and cell-based assays. MV co-supervised the project. FF supervised the project and provided conceptual guidance and funding. All authors discussed the results and approved the final manuscript.

